# Isolation of cancer stem cells by selection for miR-302 expressing cells

**DOI:** 10.1101/427732

**Authors:** Karim Rahimi, Annette C. Füchtbauer, Fardin Fathi, Seyed Javad Mowla, Ernst-Martin Füchtbauer

## Abstract

Cancer stem cells receive increasing interest because they are believed to be a major reason for long-term therapy failure. The reason for the therapy resistance of cancer stem cells lies partially in their multi-drug resistance and partially in the ability to rest mitotically inactive in the hypoxic center of tumors. Due to their variable number and their often low proliferation rate, cancer stem cells are difficult to purify in decent quantities and to grow in cell culture systems, where they are easily outcompeted by faster growing more ‘differentiated’, i.e. less stem cell-like tumor cells. Here we present a proof of principle study based on the idea to select cancer stem cells by means of the expression of a stem cell-specific gene. We inserted a selectable *egfp-neo* coding sequence in the last exon of the non-coding murine *miR-302* host gene. As a stem cell specific regulatory element, we used 2.1 kb of the genomic region immediately upstream of the *miR-302* host gene transcription start. Stable transgenic CJ7 embryonic stem cells were used to induce teratomas. After three weeks, tumors were removed for analysis and primary cultures were established. Stem-like cells were selected from these culture based on G418 selection. When the selection was removed, stem cell morphology and *miR-302* expression were rapidly lost, indicating that it were not the original ES cells that have been isolated. In conclusion, we show the possibility to use drug resistance expressed from a regulatory sequence of a stem cell-specific marker, to isolate and propagate cancer stem cells that otherwise might be hidden in the majority of tumor cells.

## Introduction

Cancer is a diverse group of diseases and many aspects of carcinogenesis and tumor progression are still poorly understood. The stochastic model assuming that all transformed tumor cells have an equal tumor-forming potential, and the cancer stem cell (CSC) model suggesting that only a small subpopulation, the CSCs, can generate new tumors are two alternative models to explain carcinogenesis (Holohan, Van Schaeybroeck, Longley, & Johnston, 2013; Song et al., 2017).

Tumor cells with stem cell characteristics were described already in 1994 (Lapidot et al., 1994). The American Association for Cancer Research has defined CSCs as a cell within a tumor which potentially can self-renew and cause the heterogeneous lineages of cancer cells that comprise a tumor (Clarke et al., 2006). This definition signaled a paradigm change because it linked tumor heterogeneity to the existence of multipotent CSCs and not to secondary mutations. CSCs are known as tumor survival cells since they fuel and support the tumor to grow and develop (Teng, Wang, Kabatas, Ulrich, & Zafonte, 2018). The importance of CSCs for cancer treatment has long been controversial however, CSCs are more and more recognized as the origin of therapy resistance and tumor relapse (Li, Tiede, Massagué, & Kang, 2007). CSCs are multi-drug resistant due to high expression of drug efflux pumps like ATP-binding protein cassette transporters (Wilson et al., 2011). In addition, they have a high level of DNA repair activity, which reduces their sensitivity to radiation (Baumann, Krause, & Hill, 2008). CSCs are often slow growing and can rest in a quiescent state. The low division rate makes them insensitive to classical cancer therapies, typically targeting fast growing cells. The quiescent state bears the risk of tumor relapse from CSCs hiding as micro-metastases for long periods. The role of CSCs is partially obscured because CSCs are not a fixed self-renewing population but rather a dynamic population, which is in a kind of equilibrium with more differentiated tumor cells. These cells are not only produced by the differentiation of CSCs but they can also de-differentiate into CSCs (Gupta et al., 2011). It is still controversial whether CSCs are the only source of new tumors, but e.g. in colorectal cancer, CSCs were shown to be responsible for tumor progression (De Robertis, Poeta, Signori, & Fazio, 2018). It is now generally recognized that these cells are responsible for a large percentage of therapeutic failures (Batlle & Clevers, 2017). Models for the contribution of CSCs to tumor growth have been developed (Enderling, 2015), however, our knowledge about the biology of CSCs is still rudimentary and builds largely on analogies to non-pathological stem cells. While a deeper understanding of CSCs would enable us to develop better therapies to target these cells, some of the stem cells properties hamper their analysis.

Most attempts to isolate or enrich CSCs utilize combinations of specific surface antigens (Medema, 2013), but transgenic approaches with reporter genes driven by stem cell-specific promoters have also been used with tumor cell lines in culture (Bauderlique-Le Roy et al., 2015). Expression of SOX10/CD133 was used to isolate CSCs from adenoid cystic carcinoma (Panaccione et al., 2017). dTomato fluorescent protein driven by the promoter of Sox2 led to isolation and characterization of human breast cancer cells (Liang et al., 2013). This strategy, with GFP driven by the *ARF* promoter, has also been used to isolate perivascular cells from the primary vitreous of the mouse eye by FACS sorting. Characterization of the harvested cells by morphology and analyzing gene expression pattern proved the efficiency of the technique (Iqbal et al., 2014). This approach should also be usable in experimental tumors in animals. However, because CSCs grow slower than the tumor cells they produce, it is still challenging to isolate and grow CSCs in culture.

Stem cells markers are gene products that are predominantly or exclusively expressed in stem cells. CSC markers might be tumor type specific and different markers have been reported for particular type of cancers (Teng et al., 2018). Cell surface markers have been widely explored as stem cell markers because they are very suitable for FACS isolation of small stem cell populations. Also, they can be combined for positive and negative sorting and thus increase specificity. Cell surface proteins CD133, CD24, and CD44, for example, are the three most common stem cells markers which are used to identify the CSCs in colon cancer (Sahlberg et al., 2014). Due to their functional relevance for “stem cell-ness”, stem cell specific transcription factors (TFs) have been widely investigated. This includes Oct4, Sox2, and Nanog, which are essential for the pluripotency of embryonic stem cells (ESCs). These factors are also critical for the progression of various malignancies and are e.g. highly expressed in nasopharyngeal carcinomas compared with non-tumorous tissues (Luo et al., 2013). Expression of Oct4, Nanog, Sox2, nucleostemin, Bmi, Zfx, Tcl1, Tbx3, Dppa4, and Esrrb was reported in bladder, colon, and prostate cancer (Atlasi et al., 2007; Amini, et al., 2014). Like TFs, microRNAs (miRNAs) are involved in many cellular processes including “stem cell-ness” and cancer. Deregulated expression of over 400 miRNAs including let-7, miR-30, miR-125b, and miR-335 in cancer stem cell like cells are reported (Bao et al., 2014). Deregulation and the effect of miRNAs expression pattern in the liver cancer and breast cancer stem cells has been investigated (Lou et al., 2018; Zhang, Xu, & Zhang, 2018). Surprisingly use of miRNAs as markers for certain cell types has so far been little employed.

*MiR-302/367* (here collectively called *miR-302s*) are a group of stem cell specific miRNAs that are processed from the intron of an RNA pol II transcribed non-coding host gene. The primary RNA in human consists of 2–3 exons. The polycistronic miRNA of *miR-302* is a cluster including *miR-302*b/c/a/d and miR-367. The *miR-302* cluster processes and produces five miRNAs including *miR-367* from the first intron of the non-coding host transcript (Barroso-delJesus et al., 2008). The primary host RNA includes two exons in mouse (Rahimi et al., 2018). The miRNAs share the same seed sequence, which is also shared with miR-290-295 in mice and miR-373 in humans. Furthermore, *miR-367* is processed from the same intron sequence but contains different seed sequence (Rosa & Brivanlou, 2009). It regulates the TGF-ß signaling pathway in human pancreatic cancer cells (Z. Zhu et al., 2015) and causes stem cell like behavior in medulloblastoma cells (Kaid et al., 2015).

The function of *miR-302s* in establishing and maintaining stem cells is documented by the observation that forced expression of individual members of the *miR-302* cluster can result in the formation of iPS cells (Kelley & Lin, 2012). MiRNAs *miR-302s* alongside *miR-200* have been reported as key regulators of stem cells behavior (Balzano et al., 2018). Furthermore, *miR-302* has been shown to enhance the stem-ness of the male germline stem cells (H. Zhu, Zheng, Wang, Tang, & Hua, 2018). Besides, expression of *miR-302s* is highly correlated with the expression of other CSC markers (Volinia et al., 2014).

The *miR-302* cluster is located in the first intron of its host gene on chromosome 4 and 3 in human and mouse respectively. In human ES cells, expression of the *miR-302* cluster is conferred by 525 bp immediately upstream of the transcription start site (Barroso-delJesus et al., 2008; Barroso-delJesus et al., 2009). In mouse, we have shown that a highly conserved regulatory sequence of at least 2.1 kb, is involved in *miR-302* gene regulation (Rahimi et al., 2018).

The aim of this proof of principle project was to utilize the expression of the stem cell specific *miR-302* host gene to isolate and select CSCs from a teratoma tumor, which was induced in mice. This strategy utilizes the expression of the non-coding exons of the *miR-302* host gene to express an *egfp-neo* fusion transcript. This reporter enables the selection of the CSCs expressing the *miR-302* gene, by means of resistance to G418. Because expression of the *egfp-neo* is coupled to expression of a stem cell specific gene, we speculated that CSCs can be kept in an undifferentiated stage until the G418 selection is relieved.

The specific advantages of the proposed strategy are that it can be used to isolate small numbers of slow growing cells because one can conditionally select for undifferentiated CSCs in culture.

Teratomas are fast growing complex benign tumors that include differentiated cells representing all three germ layers (Gonzalez-Crussi & Armed Forces Institute of Pathology (U.S.), 1982). They can be used for stem cells and developmental biology research purposes (Aleckovic & Simón, 2008). Teratomas can be induced by implantation of pluripotent stem cells like ES cells e.g under the skin of immunoincompetent or isogenic mice (Lensch et al., 2007).

As a proof of principle study, we inserted a selectable reporter gene cassette (*egfp-neo*) in the non-coding stem-cell-specific *miR-302* host gene to isolate CSCs from a teratoma tumor model. The transcriptional control of the selectable reporter by a stem cell specific gene enabled us to conditionally select undifferentiated CSCs as long as the selection (G418) is present.

## Results

### *Egfp-neo* expression driven by murine *miR-302* proximal promoter in transgenic CJ7 murine ESCs

In order to select transfected ES cells expressing the murine *miR-302* host gene, an *egfp-neo* fusion cassette was made in which the *egfp-neo* was driven by the miR-302 core promoter (figure S3A). This predicted *miR-302* upstream regulatory sequence included the region from -595 to 45 bp of the first exon. CJ7 ESCs were electroporated with a linearized construct to promote random integration. G418 resistant colonies were picked and individually analyzed. About 80% of the colonies showed a weak but clear EGFP signal. To confirm that the expression of *miR-302s* is stem cell specific, we grew the selected clones in differentiation medium without LIF and feeder cells. Under such conditions, the EGFP signal was almost lost within five days (figure S1).

To further analyzing EGFP expression from the *miR-302* promoter, we picked 228 primary colonies and grew them under ES cell condition with LIF and feeder cells as well as in ‘differentiation’ medium without LIF and without feeder cells. Of these 228 clones, 118 survived under ES cell condition and 100 under differentiation condition. After five days under these culture conditions, 93% of the undifferentiated colonies showed partial to complete EGFP expression compared to 35 % under differentiation condition (table S1), confirming our hypothesis that expression of the *mmiR-302* reporter depends on the differentiation status of the cells.

In a few clones, significant EGFP expression was found after five days under differentiation conditions (figure S2). While this could be due to enhancer activity flanking the integration site, the microscopic image of the green colonies indicated a reappearing of undifferentiated cells as is frequently seen in ES cell cultures under these conditions.

### Murine *miR-302* host RNA can support coding sequence

Murine *miR-302* host RNA is a non-coding transcript of unknown function. The RNA is spliced, poly-adenylated and exported from the nucleus (Rahimi et al., 2018) and thus might be used to introduce a coding sequence to be translated. In order to test this, we introduced the fused coding sequences of *egfp-neo* into the second exon of the *miR-302* host RNA under the control of the PGK promoter (figure S3C). Transfected CJ7 cells were selected using G418. 12 randomly selected clones were analyzed by RT-PCR (figure S4) and fluorescent microscopy (figure S5). RT-PCR results showed expression and correct splicing of the *egfp-neo* containing murine *miR-302* host RNA.

### *Egfp-neo* expression driven by murine an extended *miR-302* promoter

After we had established that the *miR-302* host gene can support translation of a reporter gene, we replaced the PGK promoter with 2,120 base pairs of upstream genomic region (figure S3D), which we had found to contain additional promoter/regulatory sequence responsible for *miR-302* host gene expression (Rahimi et al., 2018). We transfected cells with this construct and selected with G418. The number of clones obtained was significantly less compared to what we had observed using the PGK promoter, confirming, that the miR-302 promoter is relatively weak (Rahimi et al., 2018). We analyzed 18 resistant clones and confirmed the presence of the transgenic cassette including *miR-302s* upstream regulatory sequence and the *egfp-neo* fusion gene.

Correct splicing of the *miR-302-egfp-neo* transcript was confirmed by RT-PCR in all 18 transgenic clones. EGFP expression was confirmed by fluorescent microscopy in the transgenic clones, even though the EGFP expression level was weaker compare to the EGFP signal driven by PGK promoter. One of these clones was selected for the generation of teratomas (figure 1).

**Figure 1:**
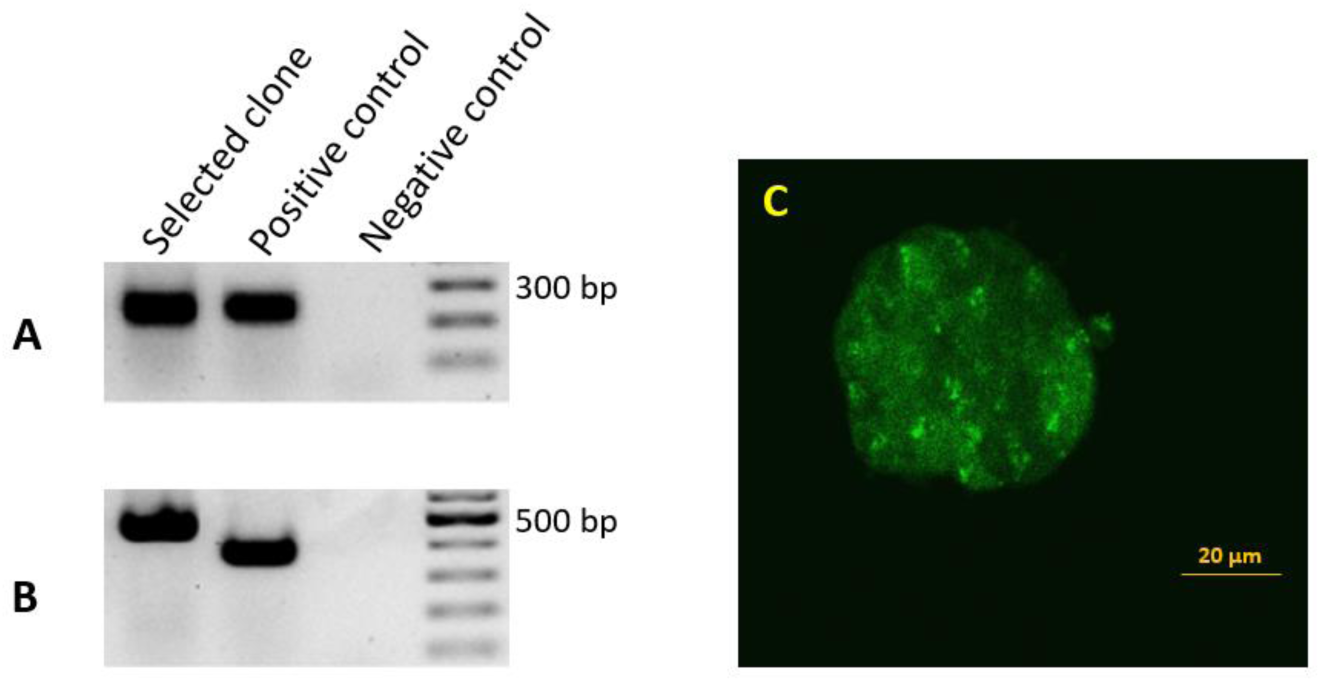
Detection of *miR-302* splicing in transgenic ES cells harboring the 2.1 kb upstream genomic region and the miR-302 host gene with *egfp-neo* in the second exon. The endogenous *miR-302* transcript results in a 221 bp band after splicing (A), while the transgene *mmiR302-egfp* transcript after splicing results in a 457 bp band (B). The positive control is cDNA from a transgenic teratoma in which exon 1 is fused to *egfp* with an expected band size of 221 bp for the endogenous *miR-302* (A) and 358 bp for *miR302-egfp* (B). Transgenic ES cells showing a weak but distinct fluorescence signal (C). Cells from this clone were used to generate teratomas.

### Primary cell culture of teratoma derived cells

In order to test our hypothesis that cancer stem cells can be selected for by the *miR-302* gene driven expression of *neo* resistance, we generated teratomas by subcutaneous injection of transgenic ES cells into isogenic mice. Three weeks after injection, when the teratomas had reached a size of around 1 cm^3,^ the mice were sacrificed and the tumors harvested and divided for primary cell culture and RNA isolation for molecular analysis.

The tumors were treated with trypsin in order to isolate single cells, which were cultured under the following four conditions: ES cell medium with and without G418 and feeder cell medium with and without G418. Cells under G418 selection were cultured on mitotically inactivated feeder cells, cells in medium without G418 were grown feeder free. Neither in ES cell nor in feeder medium, the primary cells survived G418 selection, indicating a low number of stem cells (figure 2A). After three weeks of culture in feeder cell medium without feeder cells and without selection, some fibroblast like mesenchymal cells appeared (figure 2B).

**Figure 2:**
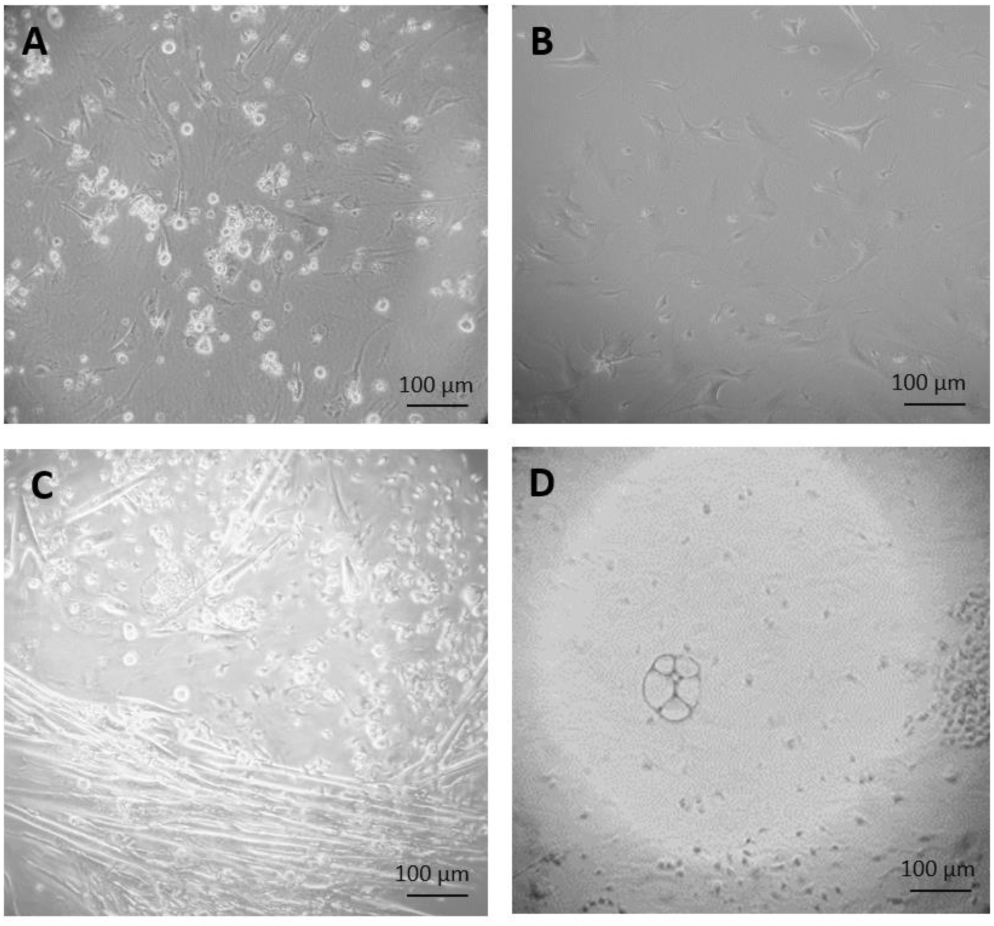
Primary culture of cells derived from teratoma. After three weeks of culture, none of the cells grown on feeder cells in ES cell medium survived selection by G418 (A). Fibroblast-like cells, after 34 days culture in feeder medium w/o G418 (B). After two weeks culture in feeder free culture in ES cell medium without G418 (C and D), some early myotubes, possibly adipocytes, and some fibroblast-like cells appeared (C). In addition, many small, rapidly growing cells appeared which were not very adhesive (D).

After two weeks of culture in ES cell medium on gelatinized dishes without feeder cells and without G418 selection, some fibroblast-like cells, some differentiated early myotubes and possible adipocytes appeared (figure 2C). We speculated that this culture might contain *mir-302* positive stem cells that we had missed in the culture, which was under immediate selection due to their low number. In order to test this, the partially differentiated culture shown in figure 2C, was split in two, one part was grown in ES cell medium and one in feeder medium, both containing G418. In this way it was possible to distinguish if the *miR-302* promoter activity was dependent on the medium condition and thus *bona fide* on the differentiation status of the cells.

In feeder medium with G418, no cells survived, indicating that cells which can grow in feeder medium do not express NEO under the control of the *miR-302* regulatory region. In contrast, if the same cells were grown in ES cell medium with G418, stem cell like colonies started to appear after two weeks while the differentiated cells disappeared (figure 3).

**Figure 3:**
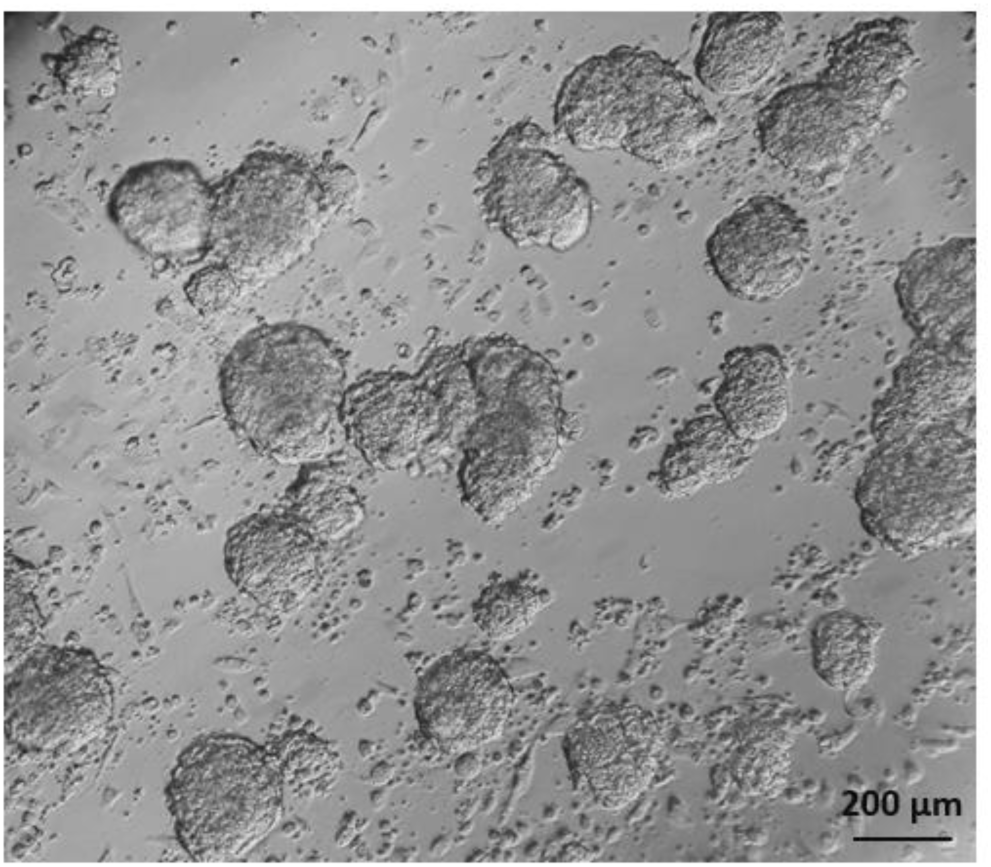
Appearance of stem cell like cells under selection. 18 days after starting G418 selection on cultures grown in ES cell medium, colonies with an ES cell like morphology were visible while the differentiated cells had disappeared.

To test if the selection for the NEO resistance coupled to the *miR-302* expression is necessary to keep the population of growing cells undifferentiated, we compared the colony morphology of cells when they were grown for three or four days after passaging either with or without G418. Without selection, the morphology of the colonies changed dramatically even though the culture conditions were otherwise identical (figure 4).

**Figure 4:**
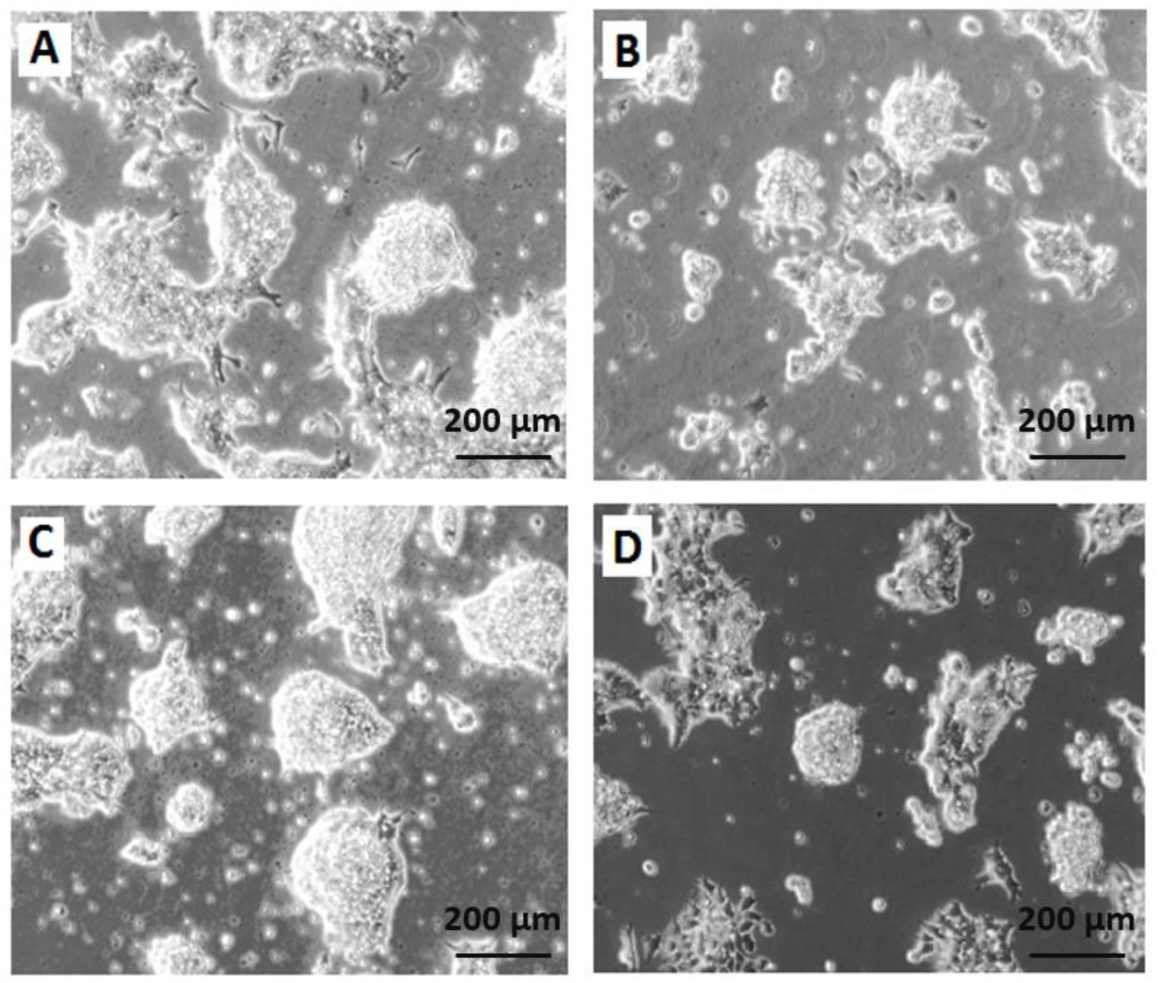
Morphology of the stem cell like colonies grown in ES cell medium with (A, C) and without (B, D) G418 for three (A, B) and four (C, D) days. Without selection, the cells rapidly lost their stem cell characteristics.

To quantify the stemness of these cells, two independent experienced persons, who did not know the culture conditions, scored colonies on 58 micrographs randomly taken from the four above-mentioned cultures (A-D in figure 4). This scoring showed clearly, that the cultures without selection were more differentiated (figure 5) than those under G418 selection. This confirmed our hypothesis that it is possible to derive and culture stem cells from tumors by coupling *neo* resistant expression to the expression of a stem cell marker like *mmiR-302*.

**Figure 5:**
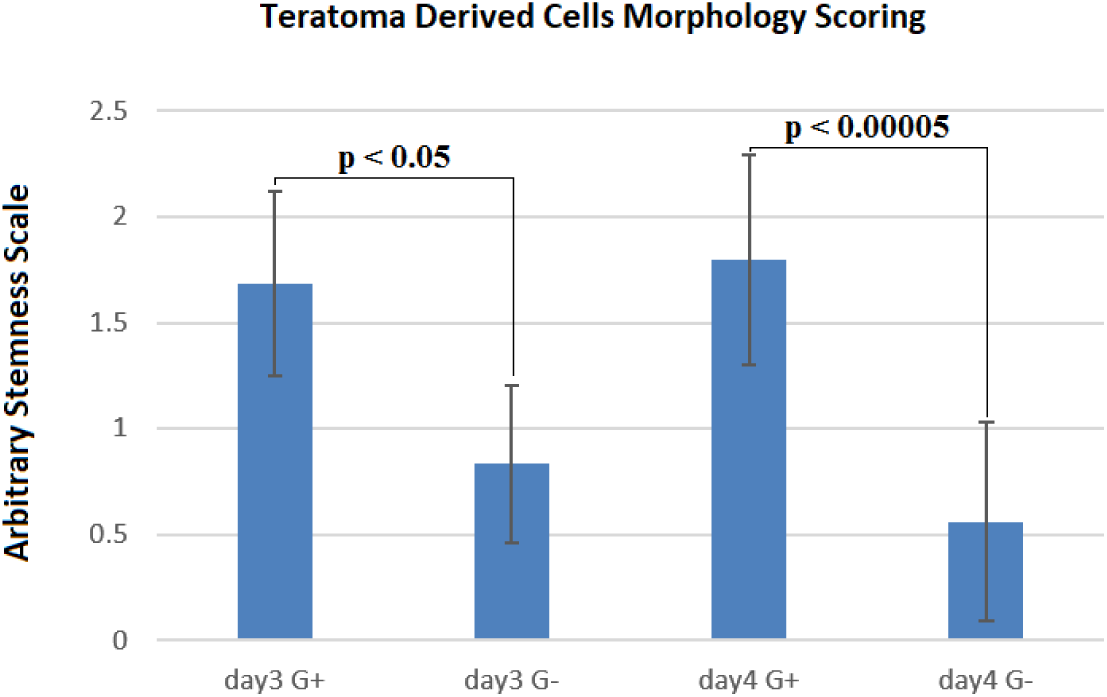
Cells under G418 selection are prevented from differentiation. On the arbitrary “stemness” scale, a completely differentiated culture would have the value 0 while a culture with all colonies undifferentiated would have a value of 3. The difference between cells with and without G418 is statistically significant after three days, but becomes more pronounced after four days. Data are shown as mean ± SD, and *p < 0.05 as significantly level. Two-tailed and unpaired t-test was performed for comparing G418 selected and unselected samples in days three and four of differentiation separately.

In order to detect the spliced *egfp-neo* RNA in the ES cell like teratoma derived cells with and without G418 selection for four days, we performed RT-PCR from exon 1 to exon 2 spanning the *egfp-neo*. This showed that the initially weak transgene expression was upregulated when the cells stably grown under selection (compare lane 1 and 2 in figure S6), but that the cells started to differentiate and down regulate *miR-302* expression when selection is released (compare lane 2 and 3 in figure S6). The short amplicon of the endogenous *miR-302* host RNA (221 bp) is under these conditions amplified beyond the exponential phase and does therefore not reflect the differences between the culture conditions.

## Discussion

CSCs are an essential factor in cancer relapse (Teng et al., 2018) and an obvious target for cancer treatment (Cianciosi et al., 2018). However, our knowledge about CSCs is limited due to the inherited difficulty to obtain pure populations of theses cells. In this proof of principle study, we show that it is possible to isolate CSCs from tumors by expressing a resistance gene under the control of a stem cell-specific regulatory element. Here we used the miR-302 host gene, including 2.1 kb of the upstream genomic region and introduced a neomycin resistance gene into the second exon. We knew that the miR-302 host RNA, which is a *bona fide* non-coding RNA is capped, spliced and exported from the nucleus (Rahimi et al 2018). We thus speculated that it might be possible to turn it into a coding mRNA. It is an interesting side aspect of our work, that this transgenic hybrid RNA can support the translation of a coding sequence. We used ES cell-derived teratoma as a well-established model tumor (Tzukerman et al., 2003). It could, therefore, be argued, that the stem cells we isolated were in fact residual ES cells and not CSCs. Two observations strongly indicate that this is not the case. The CSCs obtained after selection grew faster than CJ7 ES cells and, importantly, without selection the cells started to differentiate under culture conditions in which ES cells will proliferate without differentiation (figures 4B and 4D). In an other study, we have tested the expression of a luciferase reporter gene under the control of the same 2.1 kb genomic region. Also in this assay, expression was lost when the selection pressure was released, further indicating that the CSCs are different from ES cells (Rahimi et al., 2018). Our results confirm that the 5’ upstream genomic region of the miR-302 host gene can drive the expression of a reporter gene in a stem cell-specific pattern. These results are in agreement with the report of Jesus et al. (2009) showing that human *miR-302* is involved in the maintainance of stem-ness and self-renewal of human ES cells, an effect that is mediated by the regulation of cyclin D1 and D2 expression, that affect the balancing of the cell cycle (Card et al., 2008; Lee et al., 2008).

It was surprising that no cells survived the G418 selection in the initial culture, even though we know that it must have contained stem cells. This is evident because we could select plenty of stemcells from the differentiated culture which was prepared by the same pool of cells. There are several reasons why the original selection failed. The primary cells from the tumor might be specifically stressed by the abrupt switch from a relatively hypoxic tumor environment to a cell culture environment and thus might be especially sensitive to the selection. Likewise might conditions like cell density etc. make the cells more vulnerable. We can therefore not exclude, that different types of tumors need different types of conditions. We are also aware of the fact that CSCs from teratomas are particularly fast growing, which might have eased the selection in our example. However, the fact that the stem cells were hidden in the differentiated culture consisting of a large number of *bona fide* post mitotic cells, shows that the CSCs were by no means overgrowing or outcompeting more differentiated cells in the culture.

## Conclusion

In order to isolate cancer stem cells from primary tumors, we performed a proof of principle study in which we selected for cancer stem cells expressing the neomycin resistance gene under the control of the stem cells specific *miR-302* host gene promoter. In this transgene, the resistance gene was inserted into the last exon of the long non-coding *miR-302* host gene, which to our knowledge is a new strategy. Our result confirm that it is possible to isolate cancer stem cells by means of their stem cell specific *miR-320* expression and that it is possible to maintain the stem cell-ness of these cells by continued selection.

## Material and methods

### Statistical data analysis

A two-tailed and unpaired t-test was used to compare the G418 selected (stem cell-like cells) and unselected cells (differentiated cells) on day three and four of differentiation separately. GraphPad Prism software version 7.00 for Windows was used to perform the statistical data analysis. Results are presented as mean ± standard deviation (SD). The value of *p < 0.05 was considered for statistically significantly level.

### DNA and RNA preparation and PCR reactions

Genomic DNA was isolated according to (Laird et al., 1991). Pfu DNA polymerase (Fermentas) or AmpliTaq Gold 360 Master Mix (ThermoFisher) were used for PCR and RT-PCR reactions. M-MLV Reverse Transcriptase (Invitrogen) kit was used for cDNA synthesis according to the manual.

### Cloning protocols

FastDigest restriction enzymes from “Thermo scientific” were used for all DNA digestions according to the manuals. T4 DNA Ligase enzyme (Thermo Scientific EL0014) was used for ligation procedures according to the manufacturer’s instruction.

### The pmmiR302pGFP-NEO vector

The murine miR302pEGFP fragment was cut from the “pmmiR302pEGFP” vector (Rahimi et al., 2018) with AseI and PaeI. This fragment included the murine miR-302 core promoter region from -595 to +45 (chr3:127,544,494–127,545,132 - UCSC Genome Browser version mm9) and the *egfp* CDS. The pUC19 (GenBank Accession No. M77789.2) vector digested with NdeI and PaeI was used as a backbone to ligate the miR302pEGFP fragment in. The resulting vector “pUC19-pmmiR302pEGFP” was digested with XhoI and Bsp119I. The DNA sequence of *egfp-neo* fusion separated by the T2A self splicing sequence was obtained from the “pUC57-pPB.TET.GFP.RFP” vector (kindly provided by Dr. Mark Denham, Aarhus University) digested with XhoI and Bsp119I. The *egfp-neo* fusion sequence was ligated in the linearized “pUC19-pmmiR302pEGFP” to give the “pmmiR302pEGFP-NEO” vector (figure S3A). This vector was linearized with ApaLI for ES cell transfection.

### The pmmiR302hostGFP-NEO vector

In order to design a vector containing the whole murine *miR-302* transcript including *egfp-neo* sequence in the second exon which is the last exon of the non-coding host RNA, the pKO Scrambler NTKV-1903 vector was used as the backbone. Using AscI, the neomycin gene cassette was removed and the vector was religated resulting in the pKO/noNEO vector. The DNA sequence of the *egfp-neo* fusion was again obtained from the “pUC57-pPB.TET.GFP.RFP” vector by Bsp119I digestion and ligated into the new pKO/noNEO vector after digestion by ClaI and dephosphorylation by alkaline phosphatase. Bsp119I and ClaI digestion result in compatible ends. The first part of the gene cassette excluding promoter, was amplified by Pfu polymerase using fwd: 5’ AAGAATACCGGTAGAACAGGACTCTTTGGG; rev: 5’ AAGAATGGATCCGGGATTTGCCTTTGTGGA primers which included AgeI and BamHI sites respectively. The length of this inserted sequence was 1,603 bp (chr3:127,545,106–127,546,708 - UCSC Genome Browser version mm9) starting from the transcription start site, spanning the intron and ending 76 bp in the second exon. The PCR product was ligated into the AgeI and BamHI sites of the vector.

The second part of the gene cassette was amplified using Pfu polymerase and by fwd: 5’ AAGAATCCGCGGCAATCCAGCTATGAGTAACA; rev: 5’ AAGAATGTCGACAGAAGGGATGAGGGAGAG primers which included SacII and SalI sites respectively. The length of this sequence was 1,756 bp (chr3:127,546,782–127,548,537 - UCSC Genome Browser version mm9) and started 148 bp downstream of the second exon. The PCR product was cloned into the SacII (not FastDigest) and SalI sites of the vector. This resulted in the “pmmiR302hostGFP-NEO” vector (figure S3B).

### PGKmmiR302hostGFP-NEO

In order to investigate the processing of the *egfp-neo* in the murine *miR-302* host RNA deriving by a common promoter, the PGK promoter was inserted upstream of the *mmiR-302* host gene transcription start site. PGK promoter was cut from a vector with HindIII and PstI and was inserted in the pUC18 vector (GenBank accession number: A02710.1) which was digested with the same enzymes. This new vector was digested with PstI and SalI and ligated to a PstI and SalI digested PCR product, which had been amplified from the “pmmiR302hostGFP-NEO” vector using fwd: 5’ AAGAATCTGCAGAACTCAGGAGTTAGGAGTAG; and rev: 5’ AAGAATGTCGACCATGTTAAAGCAGAGGGGA primers. By digesting the “pmmiR302hostGFP-NEO” vector with NheI and SmaI, 3 different fragments were achieved. The 3,757 bp size fragment (including *miR-302* transcript + *egfp-neo*) was ligated to the backbone of the new “pUC18-PGKp” vector after digestion with NheI and SmaI. The obtained vector “PGKmm302hostGFP-NEO” (figure S3C) was linearized with SspI and used for the ES cell transfections.

### The pmmiR302phostGFP-NEO vector

Later 2.1 kb of the *mmiR-302* promoter region was inserted upstream of the transcription start site in the “pmmiR302hostGFP-NEO” vector. A 2,428 bp PCR amplicon including 2,120 bp of the *miR-302* regulatory sequence down to intron 1 of the miR-302 host gene (chr3:127,542,969–127,545,378 – UCSC Genome Browser version mm9) was amplified using forward primer 5’ AAGAATACCGGTCTGGAGTTGCTTTGTTTTC including an AgeI site and reverse primer 5’ AAGAATCATGTTAAAGCAGAGGGGA including a SpeI site. Digestion of the 2,428 bp PCR product with AgeI and SpeI gave two fragments of 2,249 and 176 bp respectively. By digestion the vector “pmmiR302hostGFP-NEO” with AgeI and SpeI, a fragment of 112 bp was removed and replaced with the 2,249 bp of the digested PCR product. This new vector, which called “pmmiR302phostGFPNEO” (figure S3D), was linearized by AseI and used for CJ7 cell transfection.

### Cell culture and electroporation

CJ7 murine ES cells (Swiatek & Gridley, 1993) were grown at 37 °C in ES cell medium [DMEM (Gibco 41965–039), 15% Fetal Calf Serum (Pan Biotech GmbH, 2602-P250915), 1000 U/ml LIF (Invitrogen PMC9484)] on mitotically inactivated feeder cells, unless otherwise mentioned. All cell culture dishes were gelatinized with 0.1% gelatin (Sigma-Aldrich G1393). Electroporation settings were 240 V and 500 µF, 6.6*10^6^ cells in 800 µl complete PBS. Electroporated cells was washed in 10 ml ES cell medium and seeded on 4*6 cm dishes, which were pre-seeded with feeder cells. For all transfections, 25 µg of vector were linearized and ethanol precipitated. Selection of the transfected cells was started 24–36 hour after electroporation. Selected colonies could be picked after 6–8 days. G418 concentration was ~350 µg/ml at a potency of 785µg/mg (Roche Diagnostics GmbH, 04727894001).

### Teratoma derived cells

The procedure of making the teratomas and achieving the teratoma derived cells is described in (Rahimi et al., 2018). Briefly, for each injection, 1600 stably transfected CJ7 ES cells with “pmmiR302phostGFPNEO” vector suspended in 50 µl Hank’s buffer were used. The cells were injected s.c bilaterally into the back of two 7 months old 129Sv/Pas isogenic male mice. After 21 days, tumors had reached a size of around 1 cm^3^ and mice were sacrificed by cervical dislocation and tumors harvested carefully. For primary cell culture, samples were prepared in 3–4 mm sample sizes, washed 3 times with cold calcium and magnesium-free PBS, immersed in 0.25% trypsin for 6 hours at 4 degrees and thereafter, after removing excess trypsin, incubated at 37 degrees for 30 minutes. Trypsin was inactivated by FBS and, 10^6^ cells were seeded per well of 12 wells plate in ES cell medium and feeder cell medium with and without G418 selection. 300 µg/ml G418 was used for NEO selection. Finally, teratoma derived ES cell-like cells were used for further analysis.

## Ethics Statement

Animal experiments were done according to the regulations of the Danish Animal Experiments Inspectorate, the legal authority under the Danish Ministry of Environment and Food.

